# Escape from X inactivation drives sex differences in gene expression

**DOI:** 10.1101/2025.08.01.668097

**Authors:** Carrie Zhu, Liaoyi Xu, Arbel Harpak

## Abstract

X chromosome inactivation (XCI) partially balances gene dosage between sexes, yet many genes are expressed from the inactive X (Xi) to a variable degree. In this study, we investigate whether variation in Xi expression among genes predicts transcriptional and phenotypic consequences of X-linked variation. We find that Xi expression levels are a strong linear predictor of female-male expression differences, suggesting that other compensatory or regulatory mechanisms play a more minor role in sex differences in X-linked gene expression. Among females, we identify traits—including BMI, estradiol, and testosterone levels—for which higher Xi expression correlates with the strength of evidence for either additive or dominance effects on the trait. We hypothesize that an underappreciated mechanism could generate dominance effects of X-linked variants on a trait—specifically when the variant influences skew in X inactivation. The work establishes Xi expression as important for understanding transcriptional sex differences and the physiological variation among females.

## Introduction

X chromosome inactivation (XCI) in females is traditionally viewed as an equalizer, bringing X-linked gene dosage to be on par with dosage in males by silencing one X chromosome ^1–3^. However, many genes “escape” this silencing and are expressed from the inactive X (Xi)^4,5^. Roughly 15–30% of X-linked genes show some degree of escape, with some escaping consistently across tissues and individuals, and others showing context-dependent escape^6,7^. The extent of escape can vary continuously, from low-level to equal expression between Xi and the active X (Xa). Understanding the consequences of expression from Xi is critical for interpreting both sex differences and individual variation among females^8–10^.

An immediate expectation is that levels of Xi expression would impact dosage balance between the sexes. Genes in the pseudoautosomal regions (PAR) of the X chromosome, which recombine with the Y chromosome, are not subject to XCI and thus maintain expression from both sex chromosomes^11,12^. Yet, even PAR genes show unequal expression between the sexes, with higher expression in males (from Xa and the Y homolog combined) reported in several tissues. This demonstrates that expression from Xi does not typically reach the full levels of the active X chromosome nor that of the Y chromosome^6,7,13^. In contrast, many non-PAR (NPX) genes have no Y homologs and exhibit higher expression in females due to unopposed Xi expression^6^.

Beyond sex differences, Xi expression may also influence phenotypic variation within females^14,15^. Heterozygote loss of function mutations provide compelling case studies. For example, such mutations in DDX3X increase the risks of developmental delay and disability^16^, and in KDM6A the risk of Kabuki syndrome^17–19^. These cases imply that Xi gene dosage can be impactful for development and disease. Further, Agarwal et al. recently showed that in a group of 17 NPX genes (those with Y homologs), the fitness cost of heterozygote loss of function mutations is higher than in any other group of genes genome-wide^20^.

Here, we test whether gene-level Xi expression predicts transcriptional and phenotypic consequences of X-linked variation. We find that, across genes, Xi expression levels quantitatively predict both (i) the magnitude of female-male expression differences and (ii) the strength of genetic associations with some physiological traits in females. We also suggest a novel hypothesis for non-additive effects of X-linked variants on a trait, specifically when the variant also influences a skew in X inactivation. Testing for dominance effects in X-linked variants, we find weak evidence for these effects being mediated by genetically encoded skews at individual loci. However, integrating signals across Xi expressed genes, we identify traits for which the strength of Xi expression predicts the significance of effects on traits; specifically, in the case of estradiol and testosterone, the strength of Xi expression predicts the significance of dominance effects.

## Results

We used RNA-seq data from the Genotype-Tissue Expression (GTEx) v8 project^21^ for 696 X-linked genes across 30 tissue types in 636 XY and 312 XX adult individuals. While recognizing that sex chromosome karyotype does not always align with biological sex, we henceforth label XX individuals as female and XY individuals as male. We quantify Xi expression using the average allele-specific expression data obtained from a three GTEx female donors whose X inactivation is non-random across cells and tissues (“skewed XCI”), i.e. the inactivated chromosome is the one inherited from the same parent in nearly all cells^6^. In contrast to females with a mosaic pattern of XCI in whom both parental X chromosomes are inactive in roughly equal proportions, skewed XCI females enable inference of Xi expression from bulk tissue data using allele-specific expression analysis. Namely, under the assumption that Xi expression is not higher than Xa expression, alleles at heterozygote sites with lower expression should typically, and consistently across tissues (**Text S1, Fig. S5**), be Xi-linked. Throughout, we estimated Xi expression for a given gene-tissue pair based on this approach using the subset of the three skewed XCI donors for which allele-specific expression data was available (**Text S1**).

### Xi expression predicts sex differences in gene expression

Across genes in the non-pseudoautosomal (NPX) region, expression from Xi is strongly correlated with sex differences in median expression level at both the individual tissue level (e.g., liver and skin in **Fig. 1A**) and when examining the per-gene medians across tissues (Pearson *r* = 0.64; *p* < 2.2e-16; **Fig. 1B**). Similarly, in the pseudoautosomal region (PAR), 12 out of 12 genes had higher expression in males (by 13.2% on average), and sex differences were significantly correlated with Xi expression (Pearson *r* = 0.78, *p*=0.0028; **Fig. 1B**). These results are in line with prior work by Tukiainen et al.^6^ and Tomofuji et al.^22^ who also report greater female expression in escape genes in the NPX region, greater male expression (including contributions from both X and Y paralogs) in the PAR1 region, and XCI beyond NPX. This result is, to our knowledge, the first illustration of the linear, continuous and tight relationship between Xi expression and subsequent sex differences across genes, and that it holds for both NPX and PAR regions. Importantly, this suggests that other compensatory or regulatory mechanisms play a more minor role in sex differences in X-linked gene expression.

**Figure 1:**
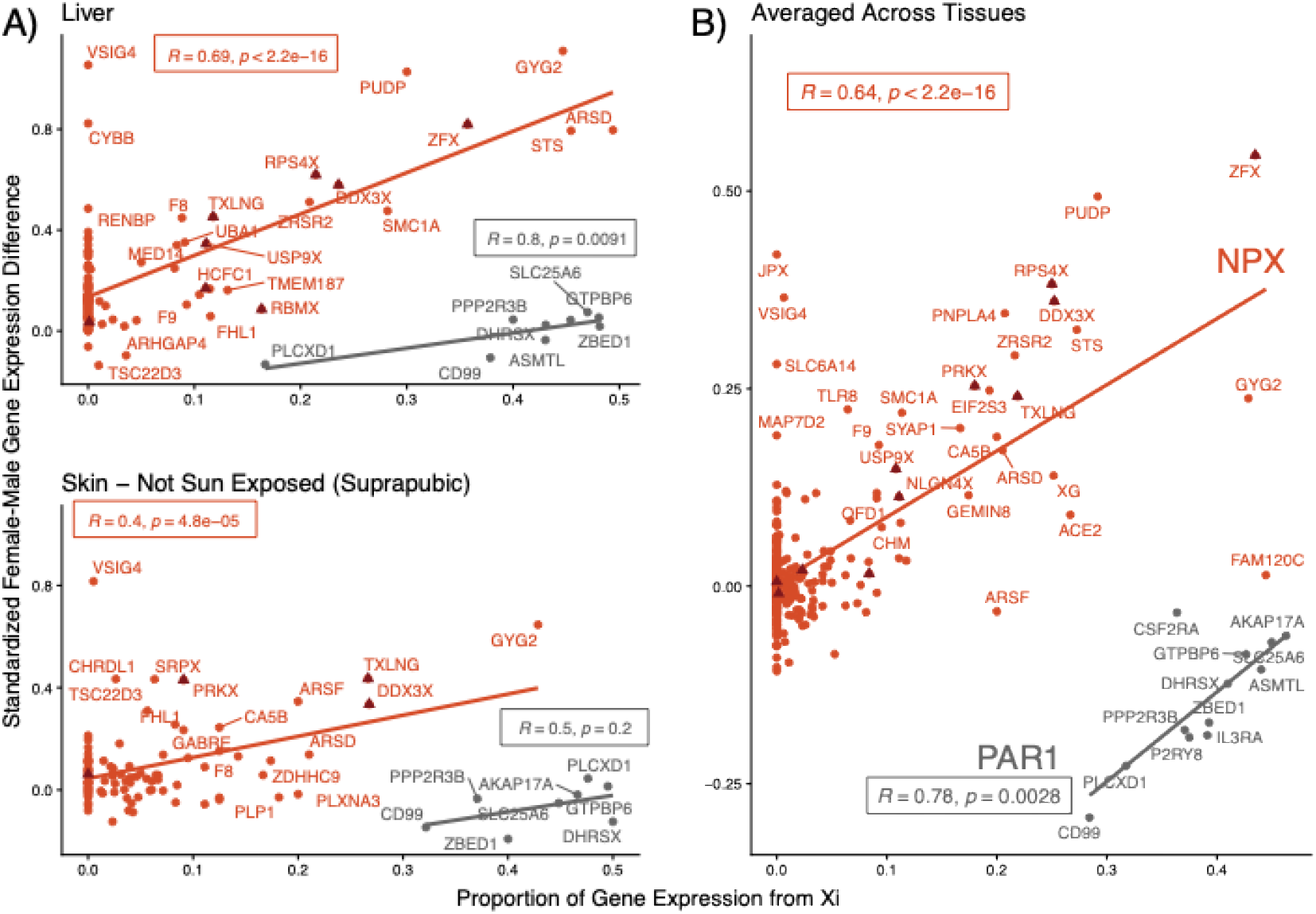
Xi expression in a single female predicts sex differences in gene expression. **(A)** The x-axis shows the ratio of expression from Xi and total expression in the tissue (liver on top, suprapubic skin on the bottom) estimated in a single GTEx female donor. The y-axis displays the difference between median female and male transcripts per million (TPM), standardized by the median male TPM for each gene. For example, y=0 confers to equal medians and 0.5 confers to the median expression of the gene among females being 50% higher than that of males. Gene expression data for the respective tissues was obtained from GTEx v8. We estimate the Pearson correlation statistics independently for the non-PAR X (orange) and PAR1 (gray) regions. Genes with homologs on the Y chromosome are indicated as dark red triangles. **(B)** The x-axis shows the median proportion of expression from Xi across tissues. The y-axis shows the median, across tissues, of standardized female-male differences.

Many Xi-expressed genes have a highly consistent expression level across tissues—18 out of 43 (42%) per a previous report^6^. In such genes, e.g. in USP11 and GDI1, we also see little variance in sex differences in gene expression (**Fig. S1**). Genes such as ARSD and FHL1, in which Xi expression is more variable, tend to also exhibit greater variance in sex differences in expression. In general, variance across tissues in sex differences is proportional to variance across tissues in Xi expression (Pearson *r* = 0.23, *p* = 0.0021 **Fig. S1**). However, we note that there are some cases the variance in sex differences across tissues is much larger than predicted by variance in Xi expression. For example, in GYG2, which is involved in glycogenesis^23^, the median expression in liver tissue is more than doubled in females compared to males, while in the esophagus’ muscularis layer it is only 20% higher in females (**Table S1**). Such tissue-specific sex differences may reflect additional regulatory factors, such as epigenetic modifications or tissue-specific chromatin states, that are not necessarily unique to Xi expression^24^.

Beyond the impact of Xi expression on sex differences, it may also contribute uniquely to female variation. We therefore also inquired into consequences for trait variation among females. Once again, we do so by examining, across genes, the relationship between Xi expression (inferred from a single individual) and measures of gene-level contribution to variation. We performed X-wide association studies (XWAS) for 21 diseases and continuous physiological traits. We chose these traits based on either having documented sex differences or relatively high SNP heritabilities for greater statistical power^25–28^. We tested for standard, additive allelic effects. We discuss the relationship between additive effects and Xi expression later in the section “**Xi expression predicts contribution to variation among females in BMI and sex hormones**. “.

### A new hypothesis for dominance effects of X-linked variants

In our XWAS, we also tested whether the heterozygote effect deviates from the average of homozygote effects (a “dominance-deviation effect”). Hypothetically, dominance may be observed on the X chromosome for the same reasons as they may exist in autosomes. However, dominance effects are thought to be negligible among common variants in autosomes^29,30^. Should XCI itself, or perhaps escape from XCI, alter our expectations for X-linked variants? We distinguish three biological scenarios for a variant in an escape gene that yield different expectations for the nature of X-linked variants’ effects.

First, we consider a biallelic SNP with an allelic effect *β* when the effect allele is expressed from Xa. If the locus harboring the SNP completely escapes X inactivation, the expected effect of a heterozygote is equal to the average of the two homozygotes (*α* = 1 in **Fig. 2-i**) just like in autosomes. When inactivation is complete and random, one of the two alleles’ effects would be zeroed at random and so the genotypic effect is halved (*α* = 0 in **Fig. 2-i**). For some intermediate degree of Xi expression (ranging from 0 to 1), the expected effect in homozygotes is the accrued effect of the allele on Xa, *β* and the effect of the allele on Xi, *αβ*— summing up to (1+*α*)*β*. In heterozygotes, the allele may be on Xa or Xi with equal probabilities (namely, ½; empty circles in **Fig. 2-i**), and so the expected genotypic effect is (filled circle for heterozygote in **Fig. 2-i**, which is on the line connecting the two homozygote effects—their average). In conclusion, the presence of a variant in an Xi-expressed locus, in and within itself, does not lead us to expect dominance-deviation allelic effects.

**Figure 2:**
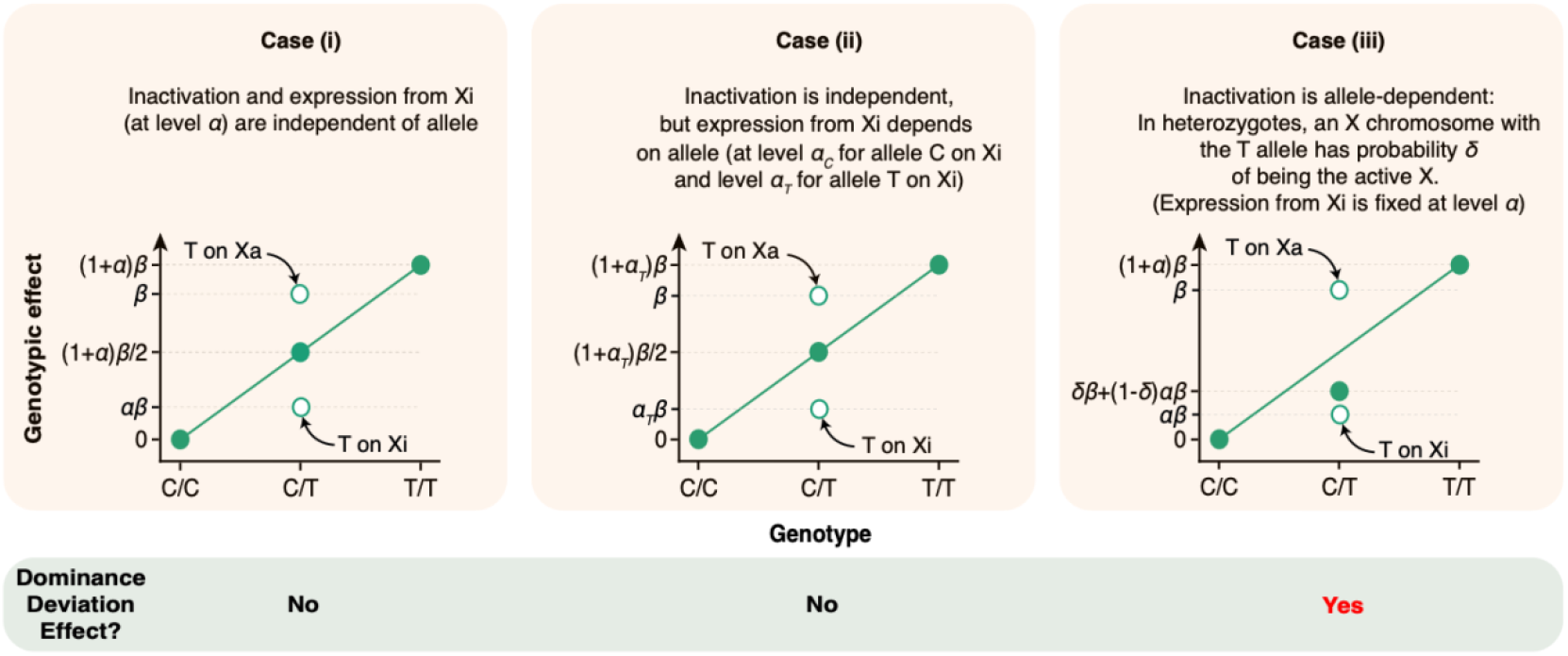
A model for the relationship between Xi expression and dominance deviation effects. The filled circles and solid lines show the expected population effects. The expected effect of heterozygotes is the weighted average of the genotypic effects in two subcases (empty circles): heterozygote individuals with the T allele on Xa and the C allele on Xi and individuals with the C allele on Xa and the T allele on Xi. In cases (i) and (ii), since inactivation is allele-independent, the weights are 0.5, and the population effect is additive. However, in case (iii), inactivation is allele-dependent, so the genotypic effect depends on the probability that a certain allele will be inactivated. If the probability is 0.5, the genotypic effect will be identical to case (i).

Second, we consider an additional effect of the variant—modulating Xi expression levels—when it is present on the inactive chromosome. Still, in this scenario, the variant does not influence X inactivation choice. While the allelic effects now depends on the allele-specific level of Xi expression, the effect remains additive (**Fig. 2-ii**).

A third case we consider is when the variant influences which parental X is inactivated. Heterozygotes will then exhibit skewing of XCI with unequal probabilities of each of the alleles being on Xa and, as a result, having the more pronounced effect. Therefore, under this scenario, we expect dominance effects in our XWAS (**Fig. 2-iii**). In other words, the allelic effect on the trait in this case is mediated by a pleiotropic effect on XCI.

Dominance effects may also arise for mechanisms that are not unique to the X or Xi specifically. We further emphasize that the mechanism proposed in case (iii) hinges on genetically encoded biases in XCI. While there is ample literature on skewed XCI in blood tissue^31–34^, the majority of skewing is thought to arise from stochastic processes, clonal expansion, or aging^34–37^, rather than from heritable genetic variation. Nonetheless, some evidence supports the possibility of heritable components to skewed XCI, especially in extreme or familial cases^38–42^. These findings may also motivate the testing for dominance effects as a way to identify potential loci that influence XCI.

Across all 21 traits, we found 117 SNPs with genome-wide independent significant (*p* < 5e-8) dominance effects in the NPX region (**Methods**) 50 of these associations were for estradiol levels, 39 for testosterone levels, 11 for glucose and only 17 for the other 18 traits. Notably, almost all of the nearest genes (69 out of 71) had no Y homologs. These effects were typically suggestive of complete dominance (**Table S2)**. To illustrate, **Fig. 3** shows the mean genotypic effect across genotypes for one SNPs with significant dominance effects on HbA1c (**Fig. 3A**), one with significant dominance effects on reported migraine cases (**Fig. 3B**), one with significant dominance effects on testosterone (**Fig. 3C**), and one with a significant effect on estradiol (**Fig. 3D**). We caveat these observations in noting that sample sizes for the minor allele homozygotes were small, and the effect of the SNPs among males (per a simple comparison of mean allelic effects conferring to hemizygote genotypes; rightmost panels in **Fig. 3**) was often discordant. One intriguing case is a SNP close to ZFP92 (**Fig. 3A**), which encodes a zinc finger protein thought to regulate DNA transcription and transposable element silencing^43^. Therefore, a linked variant may influence X-inactivation, consistent with Case (iii) in **Fig. 2**. Another intriguing case is the SNP rs147326498 which locate in the exome of MID1 (**Fig. 3C**), the protein encoded by this gene is a member of the tripartite motif (TRIM) family, also known as the ‘RING-B box-coiled coil’ (RBCC) subgroup of RING finger proteins, this gene was also the first example of a gene subject to X inactivation in humans while escaping it in mice^44^.

**Figure 3:**
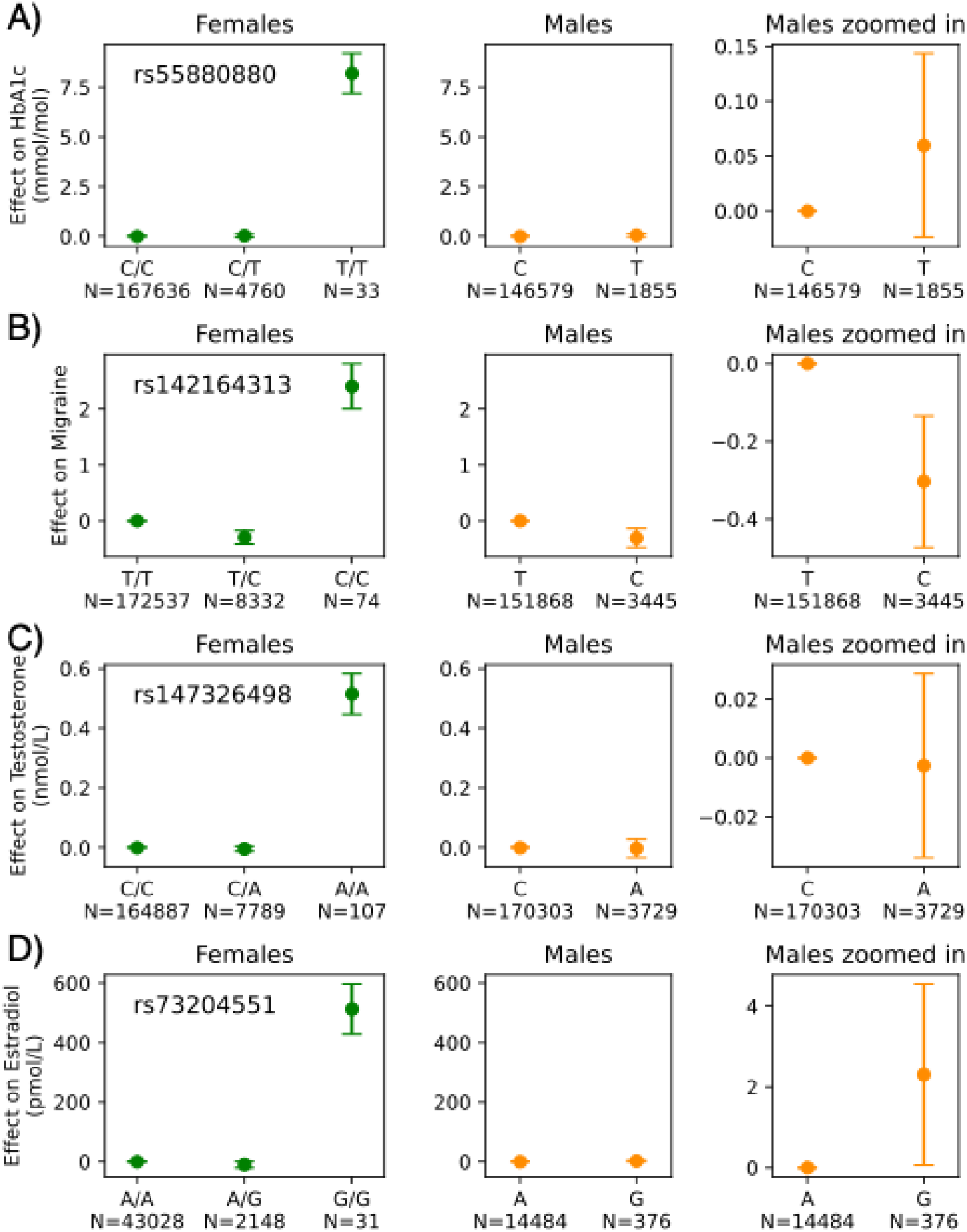
Examples of NPX SNPs with significant dominance effects. Shown are four SNPs that exhibited genome-wide significant dominance deviation effects in an XWAS for HbA1c (**A**), reported case of migraine (**B**), testosterone (**C**), and estradiol (**D**). The y-values confer to the mean allelic effects (±SE), for individuals in the White British subset of the UKB with the respective genotype. Sample sizes (N) are printed below each genotype.

### Xi expression predicts contribution to variation among females in BMI and sex hormones

In the section “**Xi expression predicts sex differences in gene expression”**, Xi expression in a handful of individuals with skewed XCI predicts sex differences in expression, which strongly suggests these Xi expression data capture levels of Xi expression in individuals with random XCI—the majority of females. With this corollary at hand, we next asked whether higher Xi expression of a gene translates to a greater role for variation among females in some traits. To address this question, we estimated the correlation between Xi expression level (using the median across tissues) and the strength of a gene’s association with the trait. To assign a strength of association, we annotated each gene with the minimal *p*-value association among SNPs for which the gene is the closest gene. We found that this measure of gene-level significance correlated with the length of the gene and/or number of SNPs associated with it (**Fig. S6, Supplementary Text S2**). We therefore reperformed this regression and included the number of associated SNPs per gene as a covariate^45^. See **Supplementary Text S3** for discussion of another approach of adjusting for the variation in the number of associated SNPs per gene^46^. We estimated this correlation across 21 traits and two types of effects, additive and dominance deviation. For BMI, cholesterol and ulcerative colitis, we found that significance of additive effects and Xi expression were highly correlated (Pearson *r* = 0.12, *p* = 0.03; *r* = 0.12, *p* = 0.028; and *r* = 0.15, *p* = 0.0063, respectively; **Fig. S3**).

If XCI skewness is heritable and more polygenic than previously appreciated, this correlation would fall in line with Case (iii) in **Fig. 2**. However, we note there is no more evidence for dominance deviation effects in X-linked variants compared with autosomes (**Fig. S7**). Overall, we find that genes with higher tissue-averaged Xi expression tend to have stronger GWAS associations near them, for both additive effects (positive Pearson correlation for 14/21 traits, 2 significant at *p* < 0.05) and dominance-deviation effects (positive Pearson correlation for 19/21 traits (1 significant at *p* < 0.05; **Fig. S3**).

We investigated this relationship further, in a tissue-specific manner, namely by examining associations of XWAS significance with tissue-specific Xi expression. Several tissue-trait-effect type associations suggest that Xi expression may influence BMI and related cardiovascular traits in females. For example, Xi expression in the artery-aorta was associated with significance of effects on BMI across genes (**Fig. 4C**)—a finding that is potentially relevant for BMI and aortic-related conditions such as arterial stiffness and atherosclerosis^47^. Similarly, Xi expression in the adrenal gland was associated with the significance of dominance deviation effects on BMI (**Fig. 4C**). Given the adrenal gland’s central role in metabolism through hormone production, including cortisol^48^, this association points to a potential mechanistic link between X-linked gene expression and metabolic regulation.

**Figure 4:**
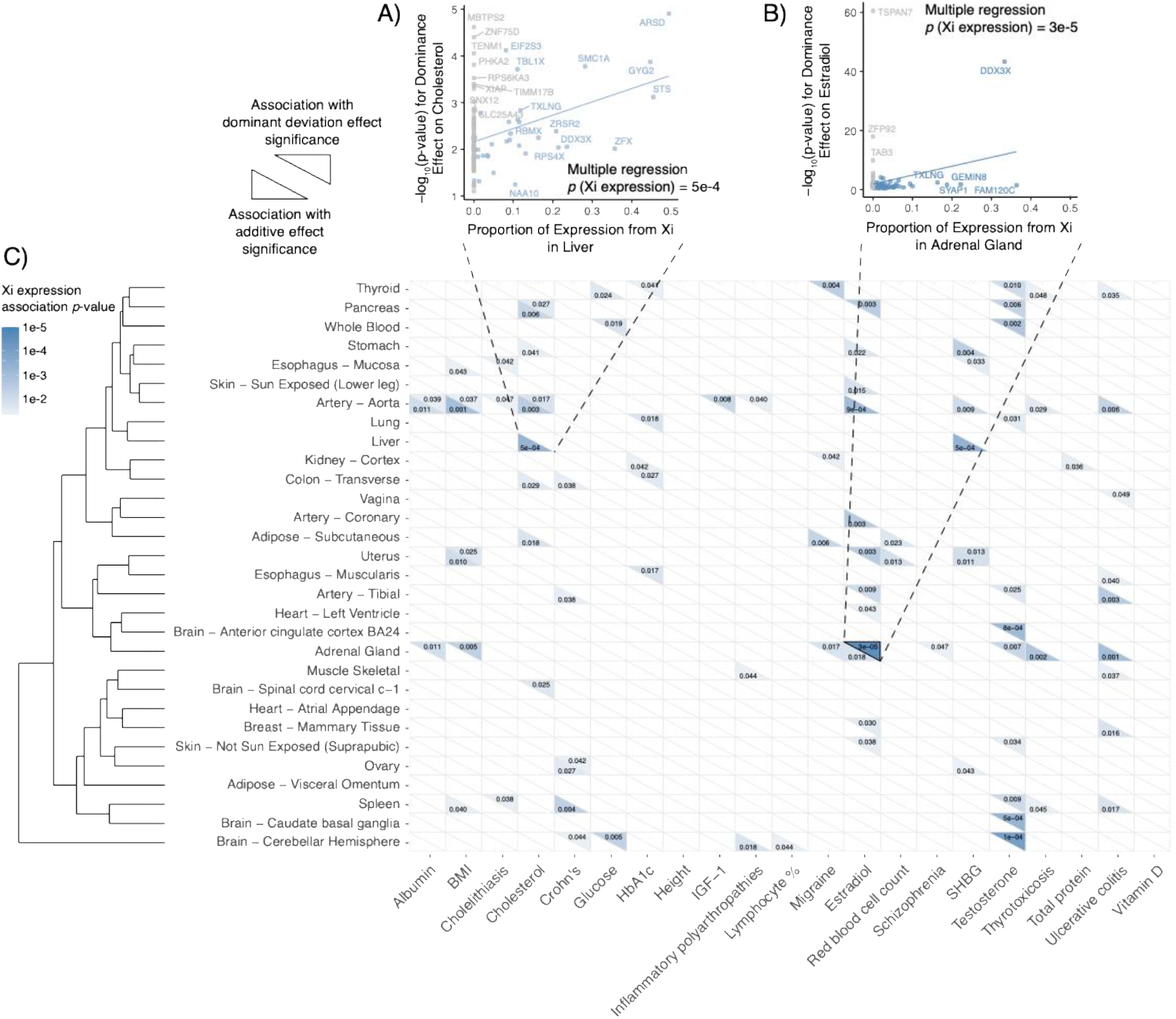
Relationship between Xi expression in a gene and its association with a trait across tissues. We estimate, for each tissue, trait and effect type (additive or dominance deviation) the effect of the proportion of Xi expression out of total (Xi + Xa) expression and the maximum -log_10_(*p*-value) among variants overlapping the gene adjusted for the number of overlapping variants. Only genes with non-zero expression (blue points) were considered. **(A)** Cholesterol in liver and additive effects. **(B)** Estradiol in adrenal gland and dominance deviation effects. **(C)** Associations across tissue, trait and effect type combinations. Only significant correlations are displayed, with blue hues conferring the significance of association. The bottom-left triangles of each block show association *p*-values for Xi expression and additive effect -log_10_(*p*-value), and the top-right triangles show the same for dominance-deviation effects. We cluster tissues by pairwise correlations of Xi expression, as shown by the dendrogram on the left. The length of each branch is inversely proportional to the correlation between tissues in Xi expression, for example, Xi expression is more correlated between lung and liver than it is between thyroid and pancreas. correction^49^ across traits, tissues and effect types; bolded top triangle circumference in **Fig. 4B**). The adrenal cortex produces androgens that can be converted into estradiol in peripheral tissues, including the aorta. This could suggest a role for Xi expression in modulating estradiol levels^50^. For both estradiol and testosterone, the association with Xi expression levels persisted in multiple tissues. Interestingly, for testosterone, it included three of the four brain regions examined (cingulate cortex BA24, *p* = 8 × 10^−4^; caudate basal ganglia, *p* = 5 × 10^−4^; cerebellar hemisphere, *p* = 1 × 10^−4^; top triangles in **Fig. 4C**).

Xi expression in the adrenal gland also correlated with significance of dominance deviation effects on testosterone and estradiol levels (the only association that remained significant after FDR

## Discussion

Our study demonstrates that expression from Xi is a major, quantifiable contributor to transcriptional sex differences and variation among females in several traits. First, we find that Xi expression, as measured in only three individuals with skewed inactivation, strongly predicts sex differences in expression across genes and tissues. While previous work has documented sex differences in expression of escape genes^6^, our findings go further by showing that this relationship extends across the full range of Xi expression, even among genes not formally categorized as ‘escapees’, supporting the view that escape from X inactivation acts along a continuum, rather than as a binary on/off phenomenon (see highly relevant discussion in ^7,51^). The strong correlation between Xi expression and higher female expression in both NPX and PAR1 regions supports a model by which sex differences arise largely from unequal dosage, rather than from differential regulation of the active X.

We also propose a simple model to delineate an X-chromosome specific expectation for dominance-deviation effects. Namely, dominance-deviation effects will arise when allele-specific skewing affects which haplotype is inactivated. By distinguishing between effects on expression given that an allele is Xi-linked and effects on XCI, we provide a clear rationale for interpreting dominance effects specific to the context of XCI. Our results do not provide any evidence for the generality of this mechanism: in fact, we do not see an overall enrichment of significant dominance deviation in X-linked variants compared to autosomes (**Fig. S7**). However, variants that do show dominance-deviation allelic effects may lead to genetically encoded XCI bias or tissue-specific gene regulation. Finally, the association we find between gene-level significance for levels of the sex hormones estradiol and testosterone and Xi expression may reflect a more polygenic role of Xi expression in sex-specific hormonal regulation, extending on previous examples such as STS, which encodes for a steroid sulfatase that is involved in regulating levels of cholesterol, estrogens, and androgens, and shows high levels of Xi expression across multiple tissues^52^.

In sum, our study shows that escape from X inactivation—treated as a quantitative, gene-level trait—largely predicts sex differences in gene expression and impacts female trait variation. Our findings underscore the importance of Xi expression in shaping sex differences and influencing genetic architecture of physiological and disease traits.

## Methods

### GTEx expression data

To estimate gene expression levels, we used bulk tissue RNA sequencing data from the Genotype-Tissue Expression (GTEx) project v8^53^, including 636 males and 312 females. Gene expression levels were measured in transcripts per million (TPM). We used a list of X-linked genes from San Roman et al.^7^ which included 712 genes on the non-PAR (NPX) region and 15 genes on the pseudo-autosomal 1 (PAR1) region of the X chromosome. We removed genes with fewer than 10 individuals with TPM greater than 1 and tissues without corresponding Xi expression data. We also removed individual-tissue-gene entries with 0 TPM.

### Use of allele-specific expression data to measure the degree of escape from XCI

We measured expression from the inactive X chromosome (Xi) using data obtained from Gylemo et al.^54^. This study used allele-specific expression data from three female donors from GTEx (UPIC, 13PLJ, and ZZPU) that were identified as having a near complete X inactivation skew, in which the same parental chromosome is inactivated across nearly all cells. For each gene, one heterozygote site with the highest RNA sequencing coverage in the largest number of tissues is chosen. The allele with the lower combined expression across tissues is labeled the inactive X (Xi) allele. XIST is the sole exception, since the Xi allele is expected to have higher expression given the gene’s role in X inactivation. Therefore, we excluded XIST from subsequent analysis.

Among the three donors, variation in allelic expression per tissue-gene pair was generally limited (**Text S1, Fig. S6**). We quantified the degree of escape using the average ratio of Xi expression to total (inactive + active) X expression across donors. In total, the combination of GTEx expression data and allele-specific expression data is available for 311 NPX and 15 PAR1 genes across 22 tissue types.

### UK Biobank

We used the UK Biobank^55,56^ for genotype-trait association. We focused on female participants, with female labeling again based on the presence of an XX karyotype. Our quality checks of sample characteristics included removing individuals with sex chromosome aneuploidy, or a self-reported sex other than female. We excluded all individuals with 3rd degree relatedness or closer, as indicated by data field 22020. We removed individuals who had withdrawn from the UK Biobank at any time before we performed the analysis of UKB data by 6/19/2024. Furthermore, we limited participants to those who self-identified as White and as British and who were also clustered in the genetic principal component space, as indicated by data field 22006. After filtering, we had a total sample of 181,012 female individuals.

### UKB Genotype Data

We used X-linked bi-allelic SNPs with INFO scores greater than 0.8 (Resource 1967 in UKB). Following a similar quality check method as used by Neale Lab^57^, we also filtered out SNPs with call rate < 0.95, Hardy-Weinberg equilibrium test *p*-value larger than 1e-10 or minor allele frequency lower than 0.001 using *plink 2.0 alpha*^58,59^.

### Trait Data

We analyzed 21 traits including 14 continuous and 7 binary disease traits. The continuous traits were chosen for their relatively high SNP heritability estimates as estimated by the Neale Lab^57^. Disease traits were classified using ICD10 coding (data field 41270 in UK Biobank) and chosen based on known sex-specific differences in disease incidence, course or prognosis.

### XWAS

We conducted female-specific association analysis for the X chromosome using *plink 2.0 alpha*^58,59^. We included birth year and the first ten principal components (data field 22009 in UK Biobank) as covariates. We applied two different models, additive and dominance-deviation, using the *--genotypic* modifier.

### Clumping

To identify independent SNPs, we performed clumping on dominance deviation XWAS summary statistics with –clump and –clump-range command in *plink 2.0 alpha* ^*58,59*^. We set the clump window size to 250 kb of upstream and downstream of each focal SNP (--clump-kb 250), selecting the SNP with the lowest *p*-value for a nonzero dominance deviation per clump.

### Genotypic effects

We used major allele homozygotes as the reference, namely by defining their “genotypic effect” as 0, with the corresponding standard error—meaningless, for the reference— also set to 0. For heterozygotes, the genotypic effect is defined as

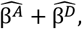

where 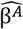 and 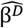 denote the estimated additive and dominance-deviation allelic effects, respectively. The standard error of heterozygote allelic effects is calculated as:

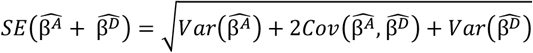

For minor homozygous, the genotypic effect is defined as

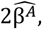

with the corresponding standard error

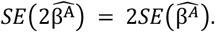

### Relationship between Xi expression in a gene and its association with a trait across tissues

To assess whether Xi expression can predict the strength of XWAS association, we considered gene level measures of significance of effect on a trait. For both additive and dominance deviation effects, we used the minimal XWAS association *p*-value among a set of SNPs in or close to the gene as a gene-level significance measure. In particular, we considered SNPs between 5000 bp upstream and 5000 bp downstream of the gene body. We regressed - log_10_(lowest SNP *p*-value) on the averaged Xi expression and the number of SNPs associated with the gene, ultimately using the *p*-value for the marginal association between gene-level Xi expression and this *p*-value as a measure of significance of association between Xi expression and strength of XWAS association. We repeated this procedure separately for the additive and dominance-deviation *p*-values for each gene.

### Summarizing the correlation distance between tissues in Xi expression using a tree

To create the dendrogram shown on the left in **Fig. 4C**, we first estimated the pairwise Pearson correlation of proportion of Xi expression levels for 14 tissues using the data from Tukiainen et al^6^. We treated the inverse of these correlations as a distance measure between each pair of tissues. Using the resulting pairwise distance matrix, we performed hierarchical clustering using the function *hclust()* in *R version 4.3.1* using default settings. The length of (horizontal) branches shown in **Fig. 4C** confers to the distance between each pair or cluster of tissues.

## Code and Data Availability

Code used in this study can be found at: https://github.com/harpak-lab/xci_sexdiff

XWAS summary statistics can be found at: https://www.harpaklab.com/data

## Acknowledgements

We thank Ipsita Agarwal, Andrés Bendesky, Jared Cole, Mark Kirkpatrick and Molly Przeworski for helpful discussions and comments on the manuscript. This work was funded by NIH grant R35GM151108 and a Pew Biomedical Scholarship to A.H. This study was conducted using the UK Biobank resource under application 92741, as approved by the University of Texas at Austin institutional review board (study protocol 00003287). The authors acknowledge the Texas Advanced Computing Center (TACC) at The University of Texas at Austin for providing computational resources that have contributed to the research results reported in this manuscript.

